# Calcium imaging and the curse of negativity

**DOI:** 10.1101/2020.09.15.298885

**Authors:** Gilles Vanwalleghem, Lena Constantin, Ethan K. Scott

## Abstract

The imaging of neuronal activity using calcium indicators has become a staple of modern neuroscience. However, without ground truths, there is a real risk of missing a significant portion of the real responses. Here, we show that a common assumption, the non-negativity of the neuronal responses as detected by calcium indicators, biases all levels of the frequently used analytical methods for these data. From the extraction of meaningful fluorescence changes to spike inference and the analysis of inferred spikes, each step risks missing real responses because of the assumption of non-negativity. We first show that negative deviations from baseline can exist in calcium imaging of neuronal activity. Then, we use simulated data to test three popular algorithms for image analysis, finding that suite2p may be the best suited to large datasets. Spike inference algorithms also showed their limitations in dealing with inhibited neurons, and new approaches may be needed to address this problem. We further suggest avoiding data analysis approaches that may ignore inhibited responses in favor of a first exploratory step to ensure that none are present. Taking these steps will ensure that inhibition, as well as excitation, is detected in calcium imaging datasets.

## 1 Introduction

The advent of Genetically Encoded Calcium Indicators (GECI) has revolutionized neurosciences by allowing the imaging of neuronal activity (Nakai, Ohkura et al. 2001, Pologruto, Yasuda et al. 2004, Tian, Hires et al. 2009), and they are now being integrated in other fields (Balaji, Bielmeier et al. 2017, Shannon, Stevens et al. 2017, Stevenson, Vanwalleghem et al. 2020). A simultaneous boom in microscopy techniques has allowed scientists to image the activity of neurons from animals such as larval zebrafish (Wyart, Del Bene et al. 2009, Ahrens, Li et al. 2012, Constantin, Poulsen et al. 2020, Vanwalleghem, Schuster et al. 2020); flies (Wang, Wong et al. 2003, Suh, Wong et al. 2004), and rodents (Chen, Cichon et al. 2012, Cai, Aharoni et al. 2016, Klioutchnikov, Wallace et al. 2020) *in-vivo* in real time thanks to GECIs. Computational tools are constantly being developed to process and extract the neuronal activity from the vast datasets that these imaging methods generate (Mukamel, Nimmerjahn et al. 2009, Freeman, Vladimirov et al. 2014, Pachitariu, Stringer et al. 2017, Giovannucci, Friedrich et al. 2019, Stringer and Pachitariu 2019). A common assumption in most of the modern computational tools is the non-negativity of the GECI’s signal.

However, negative deviations from the fluorescence baselines have been observed, and these assumptions of non-negativity may bias the results and observation by excluding relevant responses that do not show the expected peaks of activity above baseline (Favre-Bulle, Vanwalleghem et al. 2018, Marquez-Legorreta, Constantin et al. 2019, Zimmerman, Huey et al. 2019). With the slow rise and decay of GECI probes, in the hundreds of milliseconds, a long-term average firing rate above 0.1Hz would be convolved as a constant fluorescence increase above baseline. Such constant activity can be found in vestibular neurons, even at rest (Shimazu and Precht 1965, Cullen and McCrea 1993), and in the primary visual cortex neurons (Baddeley, Abbott et al. 1997) among a great many others.

Inhibition of tonically active neurons has been observed with electrophysiology in vestibular neurons(Shimazu and Precht 1966), Purkinje cells (Tian, Tep et al. 2013), or distributed across the brain in response to stimulus-driven decisions (Steinmetz, Zatka-Haas et al. 2019). Inhibition of tonic neurons, convolved by the slow GECI kernels, would translate to negative deviations from baseline as we and others have observed (Favre-Bulle, Vanwalleghem et al. 2018, Zimmerman, Huey et al. 2019).

Many tools for GECI analysis include methods for inferring the spike train that generated the observed fluorescence signal, and again most of these spike deconvolution algorithms assume non-negativity (Vogelstein, Packer et al. 2010, Pachitariu, Stringer et al. 2018). The spikefinder online challenge had this implicit assumption in the datasets offered to the community (Theis, Berens et al. 2016), meaning that the best performing algorithms were based on convolutional neural networks. This supervised approach, however, would miss response profiles that were absent in their training datasets as a result of the assumption of non-negativity.

Finally, we can find this non-negative assumption present during the analysis of the neuronal responses extracted from the calcium datasets. For example, Non-negative Matrix Factorization (NMF) has been used as a dimensionality reduction or clustering tool (Freeman, Vladimirov et al. 2014, Mu, Bennett et al. 2019, Torigoe, Islam et al. 2019). Another approach that we and others have used is the binarization of the data based on a threshold of activity to generate “bar codes” of the brain activity, which also has an intrinsic non-negative assumption (Kubo, Hablitzel et al. 2014, Naumann, Fitzgerald et al. 2016, Heap, Vanwalleghem et al. 2018, Daviu, Fuzesi et al. 2020, Etter, Manseau et al. 2020). Other threshold-based approaches, or even data cleaning steps, could also discard all the negative deviations from baseline, biasing conclusions drawn from the dataset to exclude inhibition from the modeled system.

We find this non-negative assumption at all levels of calcium imaging analysis, from the fluorescence extraction, to spike inference and neuronal response analysis. Our goal here has been to assess how the most popular calcium imaging analysis toolboxes deal with negative deviations from the baseline, which we assume come from inhibited tonic neurons. We also hope to spark a discussion on how these assumptions may have biased past studies, and may continue to bias future studies using GECIs.

## 2 Materials and Methods

The imaging data came from (Favre-Bulle, Vanwalleghem et al. 2018). Briefly, experiments were carried on 6 day post-fertilization (dpf) *nacre* mutant zebrafish (*Danio rerio*) larvae of the Tuple Longfin strain carrying the transgene *elavl3:H2B-GCaMP6s* (Chen, Wardill et al. 2013). The larvae were immobilized in 2% low melting point agarose (Progen Biosciences, Australia) and imaged using a diffuse digitally scanned light-sheet microscope (Taylor, Vanwalleghem et al. 2018) while an optical trap was applied to the otolith to simulate acceleration (Favre-Bulle, Stilgoe et al. 2017, Favre-Bulle, Vanwalleghem et al. 2018, Favre-Bulle, Stilgoe et al. 2019, Favre-Bulle, Taylor et al. 2020). All procedures were performed with approval from the University of Queensland Animal Welfare Unit in accordance with approval SBMS/378/16/ARC.

Artificial datasets were generated using the Neural Anatomy and Optical Microscopy simulation toolbox (Charles, Song et al. 2019). We used the parameters for nuclear simulation with GCaMP6f default (see table 1). To simulate inhibited neurons responses, we randomly attributed a spike number from a Poisson distribution (λ of 1, based on (Baddeley, Abbott et al. 1997)) to each 200ms time window of ten to twenty percent of all simulated neurons (since ~20% of neurons were inhibited when observed by (Steinmetz, Zatka-Haas et al. 2019)). We then set a time frame of 0.2 to 5 seconds of inhibition (0 spikes), which was used to simulate the neuronal activity and generate movies that were processed with the tools below.

**Table 1:**
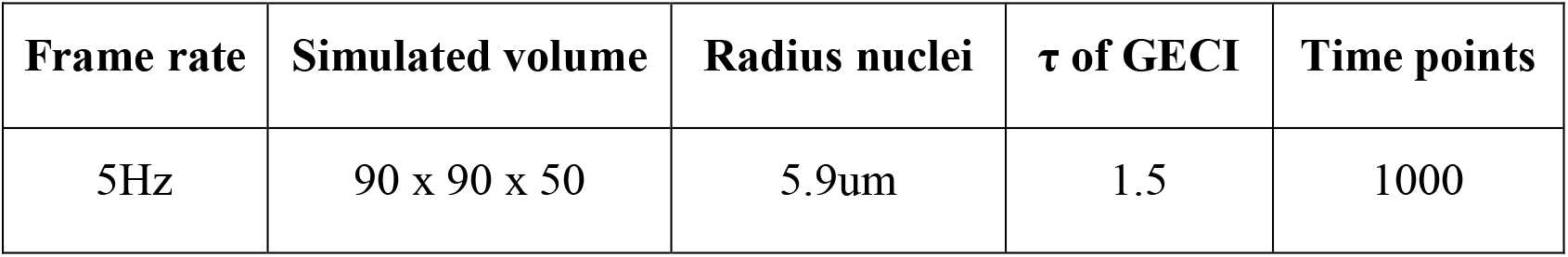
Parameters used for the simulation of calcium datasets.

For fluorescence extraction and spike inference, we benchmarked the most cited calcium imaging toolboxes: suite2p (suite2p, version 0.8.0, RRID:SCR_016434) (Pachitariu, Stringer et al. 2017), CaImAn version 1.8 (Giovannucci, Friedrich et al. 2019), and the PCA/ICA approach CellSort (Mukamel, Nimmerjahn et al. 2009). We did not simulate motion, and as such did not use the registration algorithms included in either suite2p or CaImAn. The parameters used for each of these approaches can be found the github repository. Briefly, for suite2p we used the sourcery roi extraction, with a τ of 2, frame rate of 5, diameter of neurons (4,6), threshold scaling of 0.5 and a high pass of 50. For CaImAn, we used the CNMFE, a τ of 2, frame rate of 5, a gSig of 4 and autoregressive order of 2. For the deep-learning spike inference method CASCADE, we used the Universal_5Hz_smoothing200ms pretrained model to infer the spikes on our dataset (Rupprecht, Carta et al. 2020).

For the analysis of the responses, we used MATLAB (R2018b, RRID: SCR_001622). ΔF/F_0_ was computed as in (Akerboom, Chen et al. 2012). We used the non-negative matrix factorization function nnmf with 15 factors to reanalyze the data from (Favre-Bulle, Vanwalleghem et al. 2018). We used the correlation coefficients tools from MATLAB to compute the 2-dimensional correlation between the ROIs and the ideal components, as well as between the traces or spikes and the ideal traces or spikes.

Statistical tests and plotting were done in Graphpad Prism (8.4.3, RRID:SCR_002798), we used ordinary ANOVA with Tukey’s multiple comparison test.

All the code used to generate and analyze the data can be found on github.com/Scott-Lab-QBI/NegativeCalciumResponses.

## 3 Results

### 3.1 Real data

First, we reanalyzed a zebrafish dataset from our previous study of vestibular processing in which we identified inhibited responses in hundreds of neurons across the thalamus and cerebellum (Favre-Bulle, Vanwalleghem et al. 2018). For this analysis, we focus on two neurons from the cerebellum and hindbrain of a larval zebrafish (Fig.1A) as larvae were subjected to vestibular stimuli (Fig.1B, shaded areas). As seen in the raw data (Fig.1B, arrows), we observe negative deviations from baseline during stimulation (Fig.1B, magenta traces), as well as positive responses (Fig.1B, green). Our first observation is that the classical ΔF/F0 approach with a moving baseline window (Akerboom, Chen et al. 2012) creates positive artefacts following negative deviation from baseline as seen in Fig.1B (arrows). These positive artefacts could be construed as actual responses by some approaches, since they peak at the same level as the actual responses (magenta traces with arrows versus adjacent green traces in Fig. 1B). In the ΔF/F0 trace, the results do not correlate as well for the (magenta) inhibited neuron (ρ=0.599) when compared to the (green) activated neuron (ρ=0.979). However, the z-scored trace is perfectly correlated to the raw trace (ρ=1) for both neurons. As such, we recommend the use of z-score as a normalization of calcium traces, and we will use this normalization in the following analysis.

**Figure 1:**
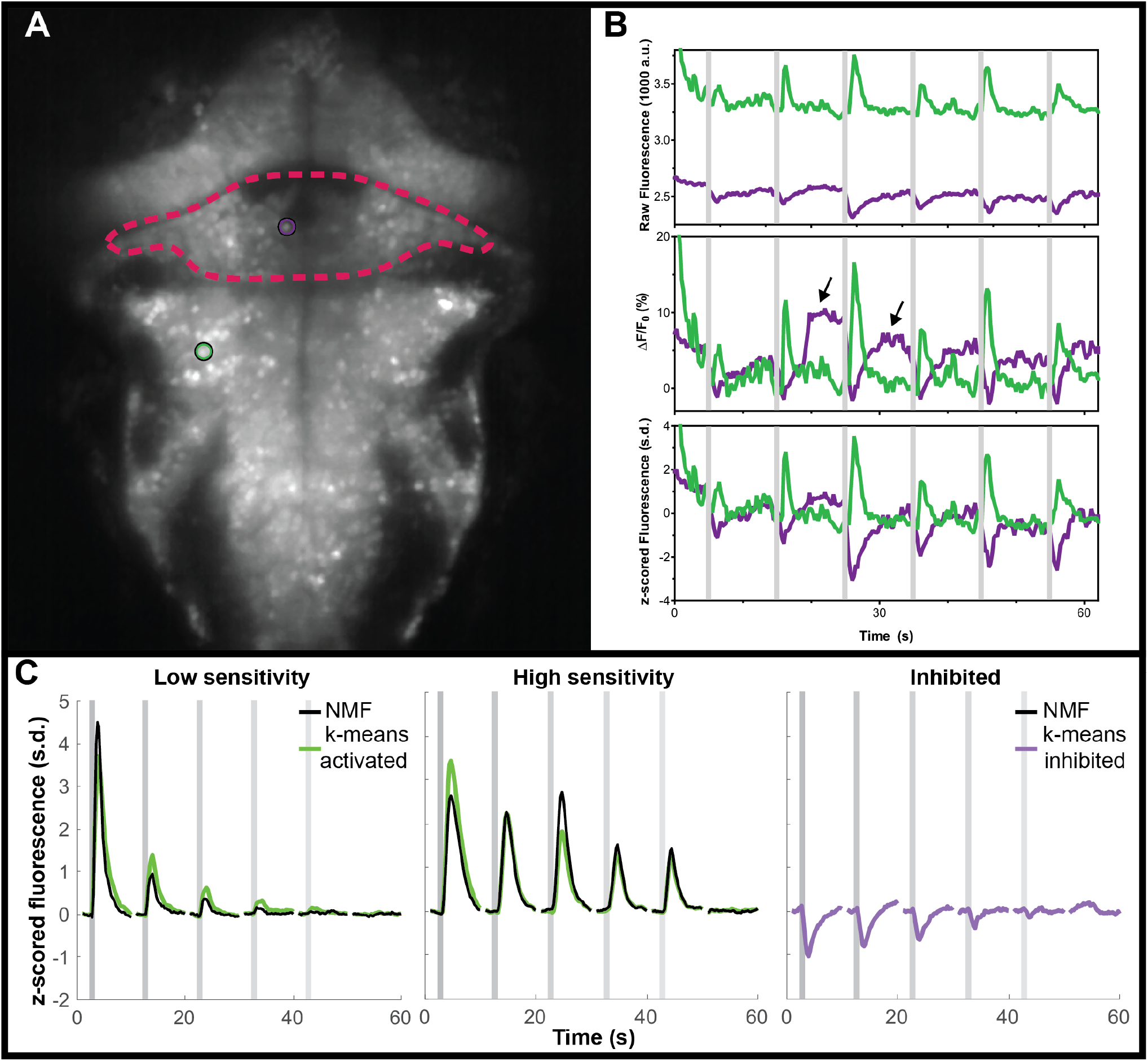
Negative deviations from baseline in real data from the cerebellum of zebrafish, and performance of various analysis tools. (A) Mean fluorescence image of a 6 dpf zebrafish expressing nuclear-targeted GCaMP6s (Chen, Wardill et al. 2013). The cerebellum is outlined in red, and am inhibited neuron is indicated with a magenta circle. The green circle indicates an activated neuron in the hindbrain. (B) Time traces of the raw (top), ΔF/F_0_ (middle) or z-scored (bottom) fluorescence for these two neurons, in their respective colors. Arrows indicate artefactual positive deviations resulting from the cessation of inhibition on the inhibited neuron. (C) Comparisons between the clusters identified using k-means (green for activated, magenta for inhibited) and those identified with NMF (black). No inhibited cluster was identified by NMF. Grey shaded areas indicate the time of vestibular stimulation (Favre-Bulle, Vanwalleghem et al. 2018), with a progression from strong to weak stimuli across the stimulus train.

An additional hurdle when working with inhibited neurons in big datasets is the risk that the data analysis methods could miss inhibited response profiles. NMF has been used to analyze larval zebrafish calcium imaging data (Mu, Bennett et al. 2019, Torigoe, Islam et al. 2019), so we tested this method on our the same vestibular dataset from our group (Favre-Bulle, Vanwalleghem et al. 2018). As can be seen (Sup.Fig.1) the NMF approach failed to identify responses resembling the inhibited cluster identified by k-means while the other (non-negative) clusters were found with a high correlation (ρ=0.92, ρ=0.94 respectively, Fig.1C).

The major limitation of this analysis is that it lacks a ground truth, making it impossible to judge whether outputs from apparently successful approaches actually reflect physiology. To solve this problem, we turned to simulated data for which we know the ground truth.

### 3.2 Simulated data

We used the Neural Anatomy and Optical Microscopy (NAOMi) Simulation toolbox (Charles, Song et al. 2019) to generate 10 datasets of simulated nuclear-targeted GCaMP6f data, as described in the Materials and Methods. Briefly, each dataset contained about 90 neurons, and for each we randomly selected either 10% or 20% of the neurons to be inhibited. For each inhibited neuron, we simulated tonic firing, based on an observed Poisson distribution (Baddeley, Abbott et al. 1997), which was randomly interrupted for 0.2 to 5s to simulate inhibition (Fig.2A). We chose a random inhibition as both suite2p and CaImAn depend on the correlation between pixels to generate the ROIs, and we wanted to make the inhibited neurons as easy to identify as possible, since most methods depend on local correlations to identify the neurons. The simulated spiking (Fig.2A) was then convolved with a GCaMP6f kernel to simulate neural activity (Fig.2B), which was then used to generate movies by NAOMi (Fig.2C). As most simulated neurons would be below the detection threshold, we used NAOMi to output the ideal responses corresponding to what would be detected with a microscope. While other algorithms occasionally identified additional neurons, the effect was marginal (<1%), so we decided to use the ideal responses as ground truth for the sake of simplicity (Charles, Song et al. 2019).

**Figure 2:**
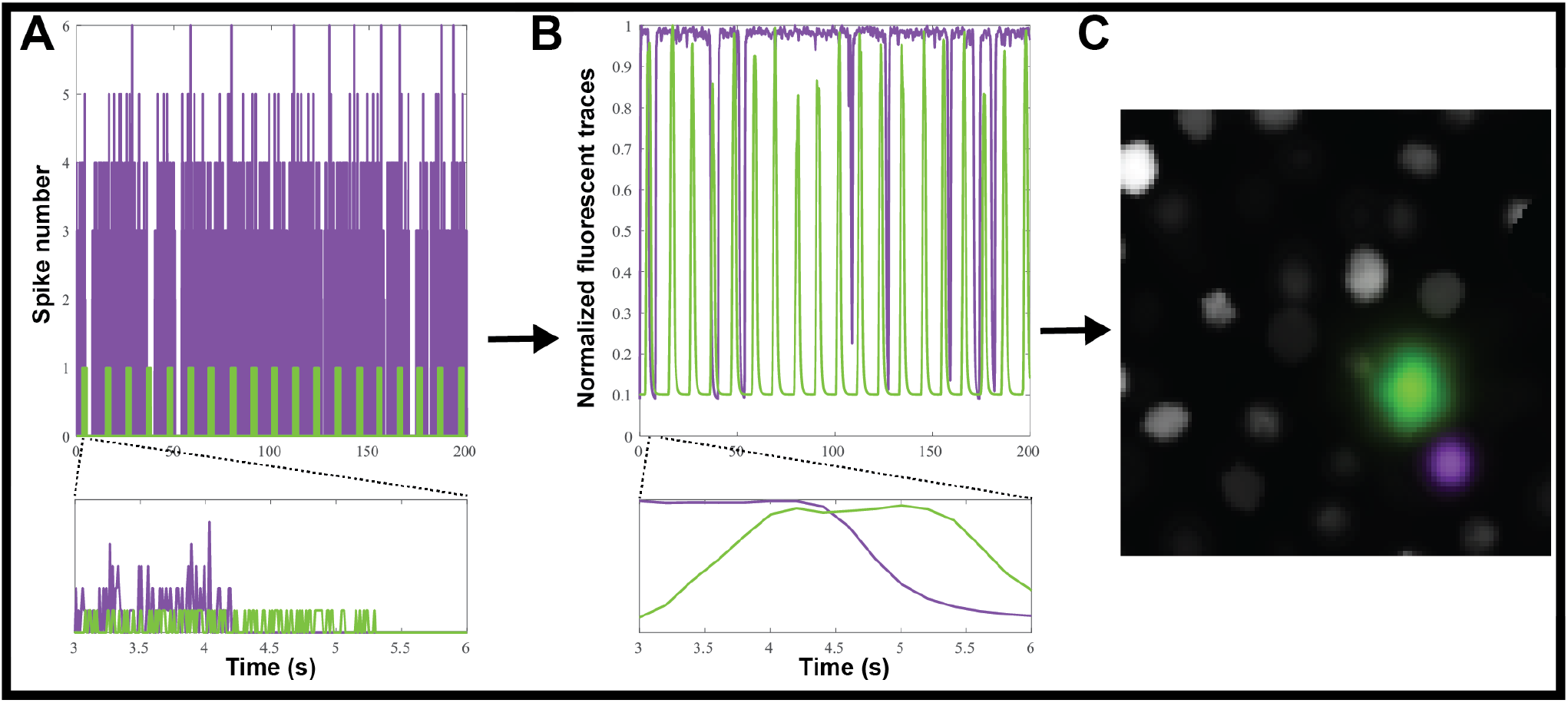
Creating simulated calcium imaging datasets. (A) An example simulated activity, showing spike numbers for one neuron (green) activated and one (magenta) inhibited by a hypothetical stimulus. (B) The spike trains are convolved with a GCaMP6f kernel and noise to generate fluorescence traces. (C) The simulated neuronal activity is used to create an artificial movie as captured by a microscope.

Each fluorescence dataset was processed through suite2p (Pachitariu, Stringer et al. 2017), CaImAn (Giovannucci, Friedrich et al. 2019), or CellSort (Mukamel, Nimmerjahn et al. 2009) and the outputs were analyzed in the exact same way. We did not investigate if either suite2p default classifier or CaImAn components evaluation would exclude inhibited neurons, as such we kept all the ROIs either algorithm identified. The raster plots of the ten datasets (Fig.3A) show that CaImAn identifies the highest number of ROIs, with CellSort and suite2p identifying a similar number of ROIs (Ideal = 94.3 ± 4.7, CaImAn = 84.7± 14.4, CellSort = 55.5 ± 3.8, suite2p = 56.1 ± 4.7).

**Figure 3:**
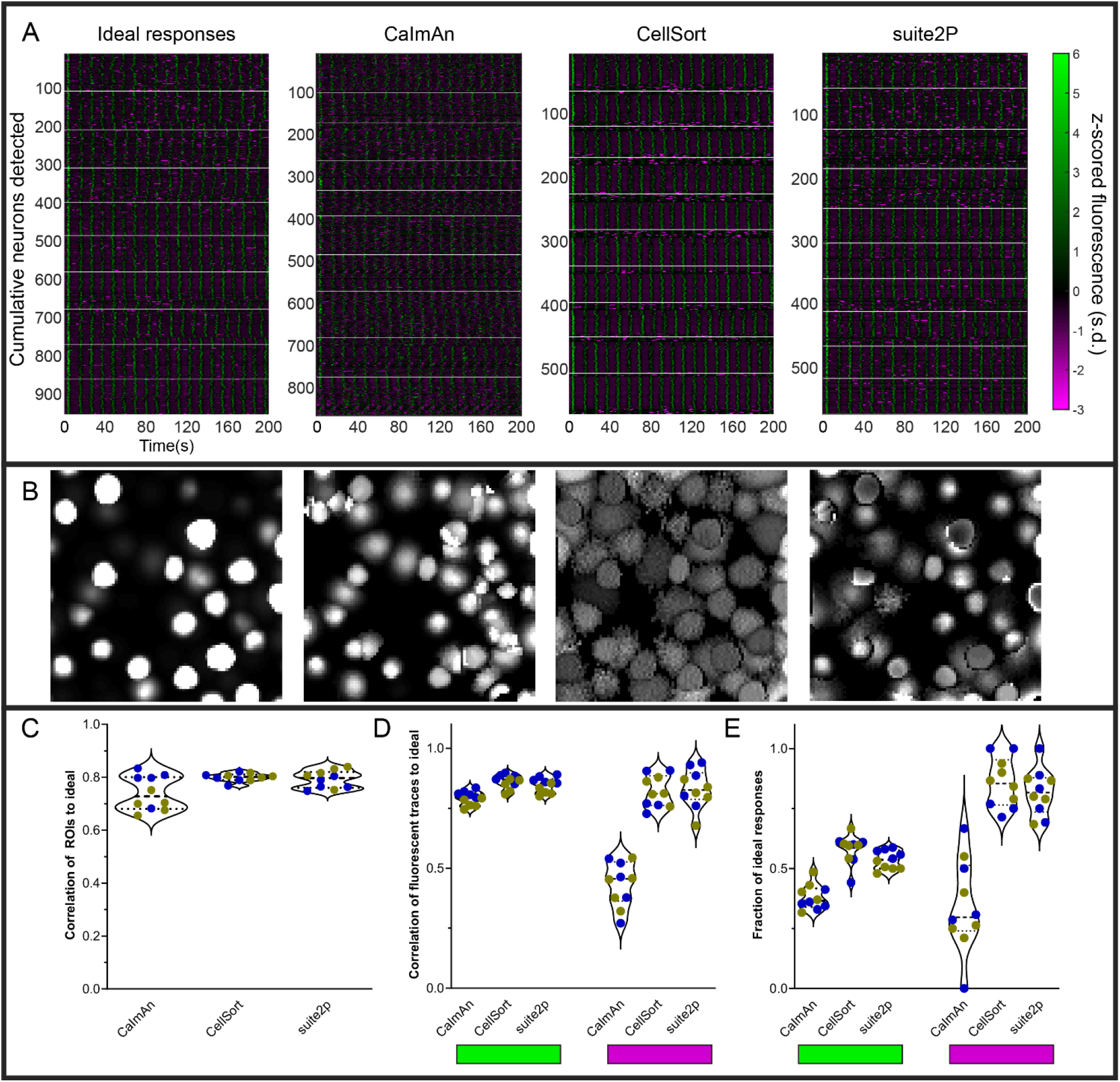
Various analyses’ performances on simulated data. (A) Raster plots of ideal responses from NAOMi, and extracted fluorescence traces from CaImAn, CellSort and suite2p. All the fluorescent traces were z-scored from −3 to 6 s.d. White horizontal lines separate the individual datasets. (B) Segmentation of the regions of interest (ROIs) by each algorithm for one representative dataset. (C) Quantification of the correlation between the ROIs identified by each of the three algorithms and the ideal ROIs. Symbol color indicate the percentage of inhibited neurons (n=5 datasets with 10% inhibited neurons in blue, and n=5 datasets with 20% inhibited neurons in yellow) (D) Average maximum correlations between the traces identified by each algorithm and the ideal responses for the activated neurons (left, green rectangle) and the inhibited neurons (right, magenta rectangle). (E) Fraction of the ideal responses identified with a correlation above 0.5 by the three algorithms for the activated neurons (left) and the inhibited neurons (right).

The segmentation of the simulated fluorescent movies gave good results for all three algorithms with well-defined regions of interest that correlated well with the ideal ROIs (Fig.3B-C, ρ=0.74±0.06, ρ=0.80±0.02, ρ=0.79±0.03). We then correlated the ideal traces of activated or inhibited simulated neurons to the traces extracted by each algorithm, for each dataset, we averaged the maximum correlations to each ideal trace (Fig.3D). We can see that all three algorithms succeeded in extracting the relevant traces for the activated neurons (Fig.3D, left, indicated by green bar), but CellSort and suite2p outperformed CaImAn for the inhibited traces (ρ=0.82±0.07, ρ=0.83±0.08, ρ=0.43±0.09 respectively, Fig.3D, right, magenta).

To assess the proportion of true positives, we identified the ideal fluorescent trace to which each ROI’s fluorescent trace best correlated. We only counted the unique ROIs that passed a 0.5 correlation cutoff, as all algorithms were shown to over-segment some of the sources in duplicated fluorescent traces (Charles, Song et al. 2019). When comparing the proportions of identified ideal activated neurons, CellSort outperformed suite2p slightly, followed by CaImAn (proportions of 0.58±0.06, 0.54±0.04 and 0.38±0.05 respectively, Fig. 2E left). For inhibited neurons, CellSort outperformed suite2p slightly again, but the divide with CaImAn grew (proportions of 0.86±0.10, 0.82±0.10 and 0.34±0.19 respectively, Fig.2E right). All algorithms seemed insensitive to the ratio of inhibited neurons present, as we saw no difference in those metrics between datasets with either 10 or 20% of inhibited neurons.

These results are lower than the results from (Charles, Song et al. 2019), who found that both CaImAn and suite2p outperformed CellSort (proportions of 0.71, 0.69 and 0.33 respectively). One possible explanation for the difference is that our use of nuclear-targeted GCaMP simulations, like our real datasets, may favor CellSort.

### 3.3 Spike inference from simulated calcium traces

In theory, inferring the spike trains responsible for the calcium traces is one way to improve the temporal resolution, as you get rid of the convolved GCaMP kernel. But the frame rate of acquisition often makes such deconvolution impractical and unreliable. Each of the above algorithms offers some form of spike inference (Fig.4A), and multiple other approaches have been proposed during an online challenge (Berens, Freeman et al. 2018). CaImAn offers multiple options for spike inference, among which we selected their fast non-negative deconvolution (FOOPSI) method (Vogelstein, Packer et al. 2010). For suite2p, we used the Online Active Set method to Infer Spikes (OASIS) (Friedrich, Zhou et al. 2017).We also tested a recent spike inference method based on deep learning, CASCADE, which offers universal pre-trained models (Rupprecht, Carta et al. 2020).

**Figure 4:**
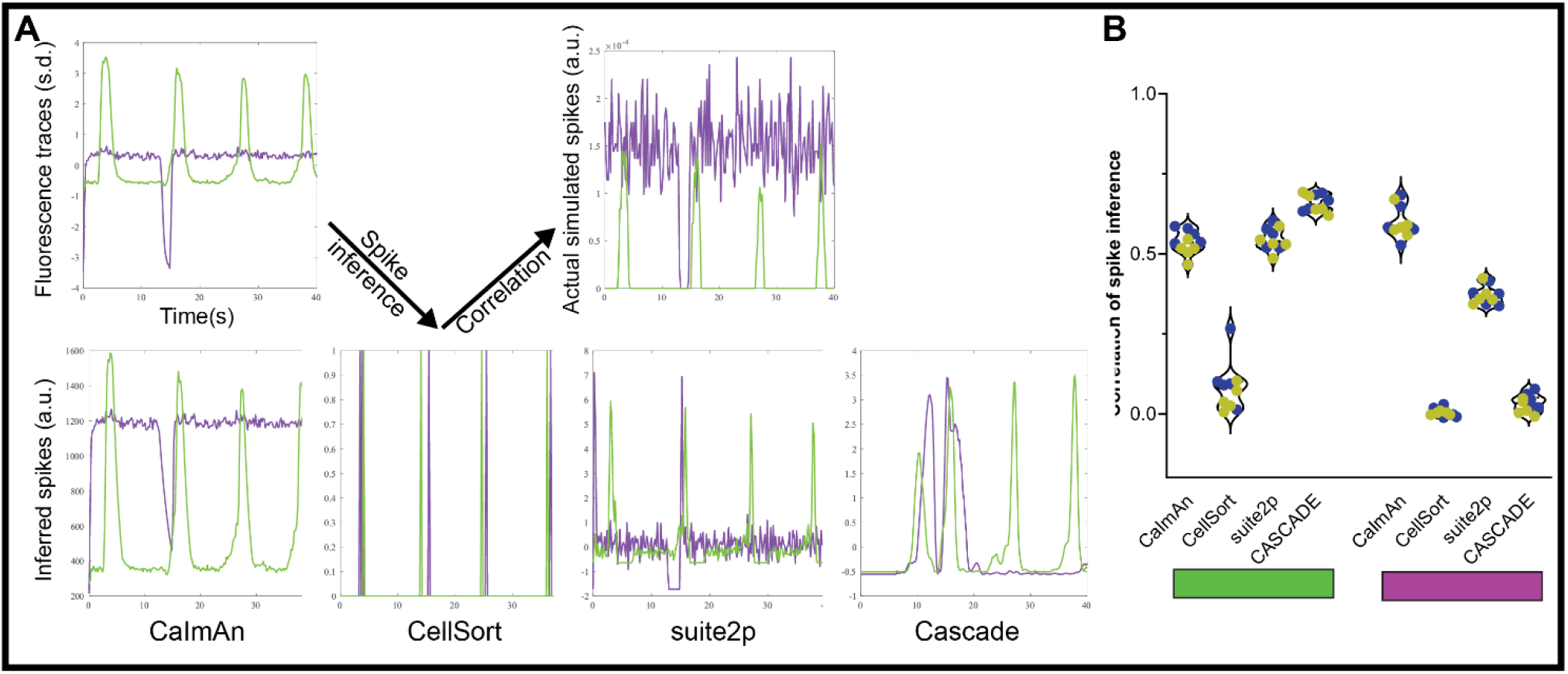
Spike inference from simulated calcium traces. (A) We used each of three methods to infer spikes from our simulated GECI fluorescence data, and compared these results to the ground truth of the actual simulated spikes used to produce our fluorescence data. (B) Correlation between the inferred spikes from the simulated calcium traces and the actual spikes for the activated neurons (left, green rectangle) and the inhibited neurons (right, magenta rectangle). Each datapoint represents the performance on one simulated dataset (n=5 datasets with 10% inhibited neurons in blue, and n=5 datasets with 20% inhibited neurons in yellow).

Using this approach, we tested how accurate each spike detection algorithm is on our datasets. To avoid any confounding issues from the detection algorithm, we used the ideal calcium responses as the basis for the spike detection. Based on our results with the moving baseline of ΔF/F_0_ (Fig.1B), we also did not pre-process the data for the spike inference with suite2p. The CellSort deconvolution approach had limited success with both activated and inhibited neurons (ρ=0.08±0.08, and ρ=0.003±0.013 respectively, Fig.4B, Sup.Fig.2). The more recent CaImAn and suite2p did well for the activated neurons, (ρ=0.53±0.04, ρ=0.55±0.03 respectively), but suite2p performance was less good for inhibited neurons (ρ=0.37±0.03) compared to CaImAn (ρ=0.60±0.05). The universal model of CASCADE performed better than the rest on the activated neurons (ρ=0.66±0.03), but worse on the inhibited neurons (ρ=0.03±0.03).

## 4 Discussion

In this study, we show that the often implicit assumption of non-negativity for calcium imaging data can lead to missing real responses from inhibited neurons. Current approaches run the risk of missing a significant fraction of responses at every step of the analysis pipeline, including cleaning the data, processing, feature extraction, dimensionality reduction, and clustering.

We have shown these negative deviations exist in real data from zebrafish, as we previously observed (Fig.1), and as observed in mice (Favre-Bulle, Vanwalleghem et al. 2018, Steinmetz, Zatka-Haas et al. 2019). We detected inhibition of tonically active vestibular neurons in the cerebellum, which may correlate to Golgi interneurons (which are responsive to tonically inhibited granule neurons) or basket and stellate interneurons (which inhibit Purkinje neurons) (Leto, Arancillo et al. 2016). Based on the dorsal position of the cell in Figure 1, they are likely stellate or basket interneurons in the superficial-most molecular layer of the cerebellum that synapse onto Purkinje neurons.

We have demonstrated that a moving baseline, such as for ΔF/F_0_, may create artefacts in inhibited neurons, which may lead to spurious generation of positive signals. Finally, inhibited responses can also be lost when using NMF or thresholding approaches to analyze and visualize the data (Fig.1C). It would be interesting to revisit the data from studies that used these approaches (Mu, Bennett et al. 2019, Torigoe, Islam et al. 2019) to see whether inhibited neurons are present in the datasets. We suggest that an initial unbiased step of data exploration of the dataset should be performed to ensure that no inhibited responses are present before pursuing steps including the above methods that assume non-negativity. Principal component analysis, or other dimensionality reduction tools, could be used to explore the data in the case of spontaneous activity or complex stimuli. Alternatively, for stimuli-driven activity, a correlation or linear regression should reveal any neuronal activity that deviates negatively from baseline.

By using simulated data (Fig.2), we tested how reliably CellSort, suite2p and CaImAn could detect inhibited neurons in a calcium imaging dataset. CellSort was the best algorithm in our specific analysis of nuclear-targeted GCaMP (Fig.3), which is at odds with other comparisons (Charles, Song et al. 2019). However, both CaImAn and suite2p are better suited to larger datasets of thousands of neurons. Between these two approaches, suite2p outperformed CaImAn for the detection of activated responses both in terms of the fidelity of the extracted response (Fig3.D, mean difference of 0.056 and p-value=0.0006) and the fraction of responses identified (Fig3.E, mean difference of 0.16 and p-value<0.0001). For inhibited responses, suite2p largely outperformed CaImAn with more than twice the fraction of ideal inhibited responses recovered (mean difference=0.47 and p-value<0.0001). Among the currently available approaches, we therefore favor suite2p, or CellSort for smaller datasets, in order to recover the most inhibited responses from calcium imaging of neuronal activity.

As for the spike inference, the algorithm included with CellSort did poorly on both activated and inhibited neurons. The more recent suite2p and CaImAn performed similarly to one another with activated neurons, in line with published results (Pachitariu, Stringer et al. 2018). However, for inhibited responses, suite2p’s performance collapsed when using OASIS. CASCADE performed well on the activated neurons, but the lack of inhibited neurons in the training datasets mean it performed poorly when detecting our inhibited responses, as such the use of a more varied training dataset could improve its performance. Overall CaImAn presents the best approach to infer spikes from inhibited neurons. Several other methods of spike inference have been benchmarked (Berens, Freeman et al. 2018), and it would be interesting to benchmark these with simulated inhibited neurons. We saw no significant differences between simulated datasets with 10% or 20% inhibited neurons in any of the above metrics, showing that the amount of inhibited neurons should not affect the detection of the activated neurons.

Overall, we suggest that the PCA/ICA approach, such as implemented in CellSort should be favored when dealing with smaller datasets and nuclear-targeted GECIs. For larger datasets however, we suggest using suite2p, which has worked well both with nuclear-targeted simulations in this study, and with a cytoplasmic GECI simulation (Charles, Song et al. 2019). When attempting spike inference, we got the best results from the FOOPSI approach, so we would favor this method when inferring spikes.

In summary, we have shown that assumptions of non-negativity can lead to the omission of real and simulated inhibited responses, and can produce spurious positive signals during the analysis of neural calcium imaging datasets. We have tested three popular and readily available approaches for analyzing such data, and provide recommendations for the best approaches to use when analyzing calcium imaging data that may contain inhibited signals.

## 7 Author Contributions

GV, LC and EKS contributed conception and design of the study; GV performed the statistical analysis; GV and LC wrote the first draft of the manuscript; GV, LC and EKS wrote sections of the manuscript. All authors contributed to manuscript revision, read, and approved the submitted version.

## 9 Acknowledgments

We thank Itia A. Favre-Bulle for data and the Vulcans who remind us “Challenge your preconceptions, or they will challenge you”. We thank C. Stringer and P. Rupprecht for helpful discussions and comments on the manuscript.

## 10 Data Availability Statement

The datasets generated and analyzed for this study can be found in the UQ espace, doi:10.14264/63584b3.

## 12 Supplementary figures

**Supplementary figure 1:**
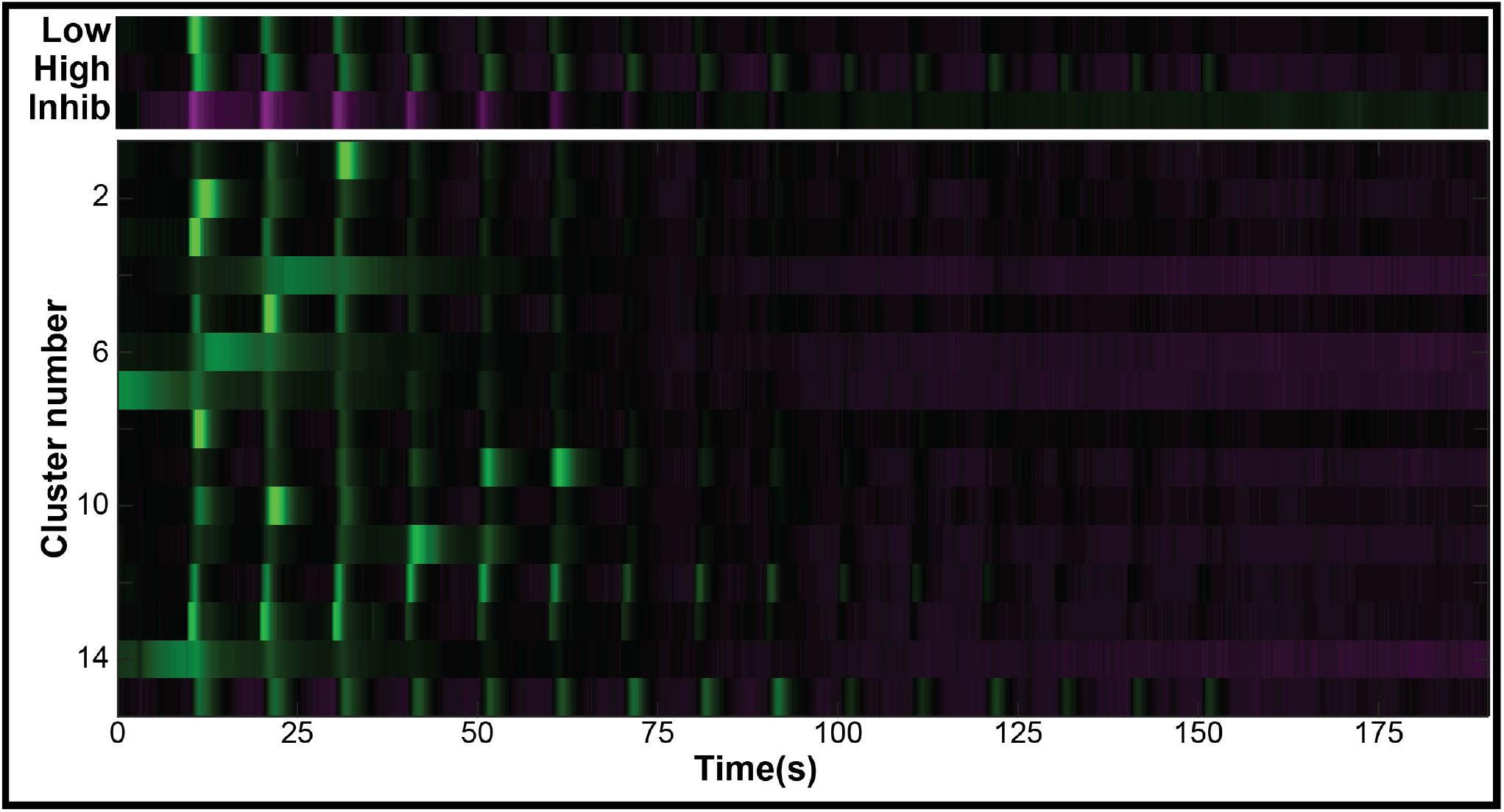
Clustering using Non-negative Matrix Factorization. The data from (Favre-Bulle, Vanwalleghem et al. 2018) were clustered with an NMF approach, using the same number of clusters as the K-means used in the original paper. Of the 15 clusters, identified the low sensitivity cluster (Cluster #3) and high sensitivity cluster (#15), but no inhibited cluster.

**Supplementary figure 2:**
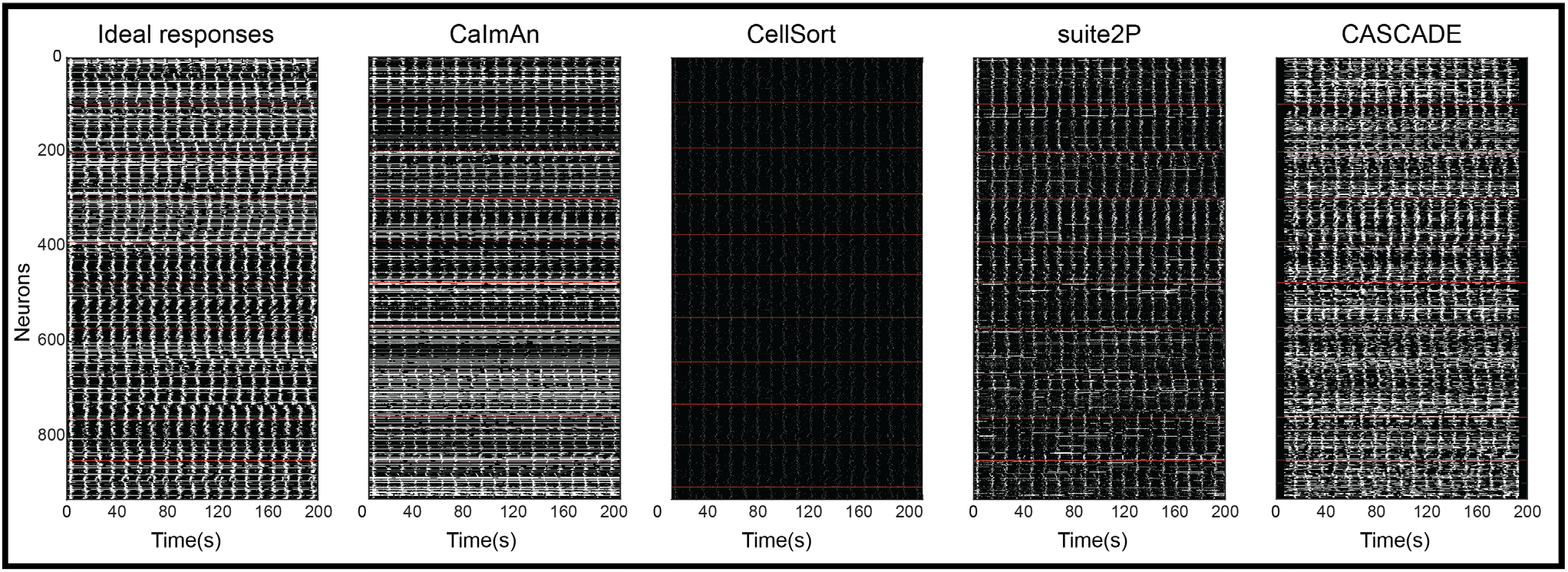
Inferred spike trains. Binarized spike trains inferred using the above algorithms from the ideal fluorescent traces.

## References

Ahrens, M. B., J. M. Li, M. B. Orger, D. N. Robson, A. F. Schier, F. Engert and R. Portugues (2012). “Brain-wide neuronal dynamics during motor adaptation in zebrafish.” Nature 485(7399): 471–477.

Akerboom, J., T. W. Chen, T. J. Wardill, L. Tian, J. S. Marvin, S. Mutlu, N. C. Calderon, F. Esposti, B. G. Borghuis, X. R. Sun, A. Gordus, M. B. Orger, R. Portugues, F. Engert, J. J. Macklin, A. Filosa, A. Aggarwal, R. A. Kerr, R. Takagi, S. Kracun, E. Shigetomi, B. S. Khakh, H. Baier, L. Lagnado, S. S. Wang, C. I. Bargmann, B. E. Kimmel, V. Jayaraman, K. Svoboda, D. S. Kim, E. R. Schreiter and L. L. Looger (2012). “Optimization of a GCaMP calcium indicator for neural activity imaging.” J Neurosci 32(40): 13819–13840.

Baddeley, R., L. F. Abbott, M. C. Booth, F. Sengpiel, T. Freeman, E. A. Wakeman and E. T. Rolls (1997). “Responses of neurons in primary and inferior temporal visual cortices to natural scenes.” Proc Biol Sci 264(1389): 1775–1783.

Balaji, R., C. Bielmeier, H. Harz, J. Bates, C. Stadler, A. Hildebrand and A. K. Classen (2017). “Calcium spikes, waves and oscillations in a large, patterned epithelial tissue.” Sci Rep 7: 42786.

Berens, P., J. Freeman, T. Deneux, N. Chenkov, T. McColgan, A. Speiser, J. H. Macke, S. C. Turaga, P. Mineault, P. Rupprecht, S. Gerhard, R. W. Friedrich, J. Friedrich, L. Paninski, M. Pachitariu, K. D. Harris, B. Bolte, T. A. Machado, D. Ringach, J. Stone, L. E. Rogerson, N. J. Sofroniew, J. Reimer, E. Froudarakis, T. Euler, M. Roman Roson, L. Theis, A. S. Tolias and M. Bethge (2018). “Community-based benchmarking improves spike rate inference from two-photon calcium imaging data.” PLoS Comput Biol 14(5): e1006157.

Cai, D. J., D. Aharoni, T. Shuman, J. Shobe, J. Biane, W. Song, B. Wei, M. Veshkini, M. La-Vu, J. Lou, S. E. Flores, I. Kim, Y. Sano, M. Zhou, K. Baumgaertel, A. Lavi, M. Kamata, M. Tuszynski, M. Mayford, P. Golshani and A. J. Silva (2016). “A shared neural ensemble links distinct contextual memories encoded close in time.” Nature 534(7605): 115–118.

Charles, A. S., A. Song, J. L. Gauthier, J. W. Pillow and D. W. Tank (2019). “Neural Anatomy and Optical Microscopy (NAOMi) Simulation for evaluating calcium imaging methods.” bioRxiv: 726174.

Chen, Q., J. Cichon, W. Wang, L. Qiu, S. J. Lee, N. R. Campbell, N. Destefino, M. J. Goard, Z. Fu, R. Yasuda, L. L. Looger, B. R. Arenkiel, W. B. Gan and G. Feng (2012). “Imaging neural activity using Thy1-GCaMP transgenic mice.” Neuron 76(2): 297–308.

Chen, T. W., T. J. Wardill, Y. Sun, S. R. Pulver, S. L. Renninger, A. Baohan, E. R. Schreiter, R. A. Kerr, M. B. Orger, V. Jayaraman, L. L. Looger, K. Svoboda and D. S. Kim (2013). “Ultrasensitive fluorescent proteins for imaging neuronal activity.” Nature 499(7458): 295–300.

Constantin, L., R. E. Poulsen, L. A. Scholz, I. A. Favre-Bulle, M. A. Taylor, B. Sun, G. J. Goodhill, G. C. Vanwalleghem and E. K. Scott (2020). “Altered brain-wide auditory networks in a zebrafish model of fragile X syndrome.” BMC Biology 18(1): 125.

Cullen, K. E. and R. A. McCrea (1993). “Firing behavior of brain stem neurons during voluntary cancellation of the horizontal vestibuloocular reflex. I. Secondary vestibular neurons.” J Neurophysiol 70(2): 828–843.

Daviu, N., T. Fuzesi, D. G. Rosenegger, N. P. Rasiah, T. L. Sterley, G. Peringod and J. S. Bains (2020). “Paraventricular nucleus CRH neurons encode stress controllability and regulate defensive behavior selection.” Nat Neurosci 23(3): 398–410.

Etter, G., F. Manseau and S. Williams (2020). “A Probabilistic Framework for Decoding Behavior From in vivo Calcium Imaging Data.” Front Neural Circuits 14: 19.

Favre-Bulle, I. A., A. B. Stilgoe, H. Rubinsztein-Dunlop and E. K. Scott (2017). “Optical trapping of otoliths drives vestibular behaviours in larval zebrafish.” Nat Commun 8(1): 630.

Favre-Bulle, I. A., A. B. Stilgoe, E. K. Scott and H. Rubinsztein-Dunlop (2019). “Optical trapping in vivo: theory, practice, and applications.” Nanophotonics 8(6): 1023–1040.

Favre-Bulle, I. A., M. A. Taylor, E. Marquez-Legorreta, G. Vanwalleghem, R. E. Poulsen, H. Rubinsztein-Dunlop and E. K. Scott (2020). “Sound generation in zebrafish with Bio-Opto-Acoustics (BOA).” bioRxiv: 2020.2006.2009.143362.

Favre-Bulle, I. A., G. Vanwalleghem, M. A. Taylor, H. Rubinsztein-Dunlop and E. K. Scott (2018). “Cellular-Resolution Imaging of Vestibular Processing across the Larval Zebrafish Brain.” Curr Biol 28(23): 3711–3722 e3713.

Freeman, J., N. Vladimirov, T. Kawashima, Y. Mu, N. J. Sofroniew, D. V. Bennett, J. Rosen, C. T. Yang, L. L. Looger and M. B. Ahrens (2014). “Mapping brain activity at scale with cluster computing.” Nat Methods 11(9): 941–950.

Friedrich, J., P. Zhou and L. Paninski (2017). “Fast online deconvolution of calcium imaging data.” PLoS Comput Biol 13(3): e1005423.

Giovannucci, A., J. Friedrich, P. Gunn, J. Kalfon, B. L. Brown, S. A. Koay, J. Taxidis, F. Najafi, J. L. Gauthier, P. Zhou, B. S. Khakh, D. W. Tank, D. B. Chklovskii and E. A. Pnevmatikakis (2019). “CaImAn an open source tool for scalable calcium imaging data analysis.” Elife 8.

Heap, L. A. L., G. Vanwalleghem, A. W. Thompson, I. A. Favre-Bulle and E. K. Scott (2018). “Luminance Changes Drive Directional Startle through a Thalamic Pathway.” Neuron 99(2): 293–301 e294.

Klioutchnikov, A., D. J. Wallace, M. H. Frosz, R. Zeltner, J. Sawinski, V. Pawlak, K. M. Voit, P. S. J. Russell and J. N. D. Kerr (2020). “Three-photon head-mounted microscope for imaging deep cortical layers in freely moving rats.” Nat Methods 17(5): 509–513.

Kubo, F., B. Hablitzel, M. Dal Maschio, W. Driever, H. Baier and A. B. Arrenberg (2014). “Functional architecture of an optic flow-responsive area that drives horizontal eye movements in zebrafish.” Neuron 81(6): 1344–1359.

Leto, K., M. Arancillo, E. B. Becker, A. Buffo, C. Chiang, B. Ding, W. B. Dobyns, I. Dusart, P. Haldipur, M. E. Hatten, M. Hoshino, A. L. Joyner, M. Kano, D. L. Kilpatrick, N. Koibuchi, S. Marino, S. Martinez, K. J. Millen, T. O. Millner, T. Miyata, E. Parmigiani, K. Schilling, G. Sekerkova, R. V. Sillitoe, C. Sotelo, N. Uesaka, A. Wefers, R. J. Wingate and R. Hawkes (2016). “Consensus Paper: Cerebellar Development.” Cerebellum 15(6): 789–828.

Marquez-Legorreta, E., L. Constantin, M. Piber, I. A. Favre-Bulle, M. A. Taylor, G. C. Vanwalleghem and E. Scott (2019). “Brain-wide visual habituation networks in wild type and &<;em&>;fmr1&<;/em&>; zebrafish.” bioRxiv: 722074.

Mu, Y., D. V. Bennett, M. Rubinov, S. Narayan, C. T. Yang, M. Tanimoto, B. D. Mensh, L. L. Looger and M. B. Ahrens (2019). “Glia Accumulate Evidence that Actions Are Futile and Suppress Unsuccessful Behavior.” Cell 178(1): 27–43 e19.

Mukamel, E. A., A. Nimmerjahn and M. J. Schnitzer (2009). “Automated analysis of cellular signals from large-scale calcium imaging data.” Neuron 63(6): 747–760.

Nakai, J., M. Ohkura and K. Imoto (2001). “A high signal-to-noise Ca(2+) probe composed of a single green fluorescent protein.” Nat Biotechnol 19(2): 137–141.

Naumann, E. A., J. E. Fitzgerald, T. W. Dunn, J. Rihel, H. Sompolinsky and F. Engert (2016). “From Whole-Brain Data to Functional Circuit Models: The Zebrafish Optomotor Response.” Cell 167(4): 947–960 e920.

Pachitariu, M., C. Stringer, M. Dipoppa, S. Schröder, L. F. Rossi, H. Dalgleish, M. Carandini and K. D. Harris (2017). “Suite2p: beyond 10,000 neurons with standard two-photon microscopy.” bioRxiv: 061507.

Pachitariu, M., C. Stringer and K. D. Harris (2018). “Robustness of Spike Deconvolution for Neuronal Calcium Imaging.” J Neurosci 38(37): 7976–7985.

Pologruto, T. A., R. Yasuda and K. Svoboda (2004). “Monitoring neural activity and [Ca2+] with genetically encoded Ca2+ indicators.” J Neurosci 24(43): 9572–9579.

Rupprecht, P., S. Carta, A. Hoffmann, M. Echizen, K. Kitamura, F. Helmchen and R. W. Friedrich (2020). “A deep learning toolbox for noise-optimized, generalized spike inference from calcium imaging data.” bioRxiv: 2020.2008.2031.272450.

Shannon, E. K., A. Stevens, W. Edrington, Y. Zhao, A. K. Jayasinghe, A. Page-McCaw and M. S. Hutson (2017). “Multiple Mechanisms Drive Calcium Signal Dynamics around Laser-Induced Epithelial Wounds.” Biophys J 113(7): 1623–1635.

Shimazu, H. and W. Precht (1965). “Tonic and kinetic responses of cat’s vestibular neurons to horizontal angular acceleration.” J Neurophysiol 28(6): 991–1013.

Shimazu, H. and W. Precht (1966). “Inhibition of central vestibular neurons from the contralateral labyrinth and its mediating pathway.” J Neurophysiol 29(3): 467–492.

Steinmetz, N. A., P. Zatka-Haas, M. Carandini and K. D. Harris (2019). “Distributed coding of choice, action and engagement across the mouse brain.” Nature 576(7786): 266–273.

Stevenson, A. J., G. Vanwalleghem, T. A. Stewart, N. D. Condon, B. Lloyd-Lewis, N. Marino, J. W. Putney, E. K. Scott, A. D. Ewing and F. M. Davis (2020). “Multiscale activity imaging in the mammary gland reveals how oxytocin enables lactation.” bioRxiv: 657510.

Stringer, C. and M. Pachitariu (2019). “Computational processing of neural recordings from calcium imaging data.” Curr Opin Neurobiol 55: 22–31.

Suh, G. S., A. M. Wong, A. C. Hergarden, J. W. Wang, A. F. Simon, S. Benzer, R. Axel and D. J. Anderson (2004). “A single population of olfactory sensory neurons mediates an innate avoidance behaviour in Drosophila.” Nature 431(7010): 854–859.

Taylor, M. A., G. C. Vanwalleghem, I. A. Favre-Bulle and E. K. Scott (2018). “Diffuse light-sheet microscopy for stripe-free calcium imaging of neural populations.” J Biophotonics 11(12): e201800088.

Theis, L., P. Berens, E. Froudarakis, J. Reimer, M. Roman Roson, T. Baden, T. Euler, A. S. Tolias and M. Bethge (2016). “Benchmarking Spike Rate Inference in Population Calcium Imaging.” Neuron 90(3): 471–482.

Tian, J., C. Tep, M. X. Zhu and S. O. Yoon (2013). “Changes in Spontaneous firing patterns of cerebellar Purkinje cells in p75 knockout mice.” Cerebellum 12(3): 300–303.

Tian, L., S. A. Hires, T. Mao, D. Huber, M. E. Chiappe, S. H. Chalasani, L. Petreanu, J. Akerboom, S. A. McKinney, E. R. Schreiter, C. I. Bargmann, V. Jayaraman, K. Svoboda and L. L. Looger (2009). “Imaging neural activity in worms, flies and mice with improved GCaMP calcium indicators.” Nat Methods 6(12): 875–881.

Torigoe, M., T. Islam, H. Kakinuma, C. C. A. Fung, T. Isomura, H. Shimazaki, T. Aoki, T. Fukai and H. Okamoto (2019). “Future state prediction errors guide active avoidance behavior by adult zebrafish.” bioRxiv: 546440.

Vanwalleghem, G., K. Schuster, M. A. Taylor, I. A. Favre-Bulle and E. K. Scott (2020). “Brain-Wide Mapping of Water Flow Perception in Zebrafish.” J Neurosci 40(21): 4130–4144.

Vogelstein, J. T., A. M. Packer, T. A. Machado, T. Sippy, B. Babadi, R. Yuste and L. Paninski (2010). “Fast nonnegative deconvolution for spike train inference from population calcium imaging.” J Neurophysiol 104(6): 3691–3704.

Wang, J. W., A. M. Wong, J. Flores, L. B. Vosshall and R. Axel (2003). “Two-photon calcium imaging reveals an odor-evoked map of activity in the fly brain.” Cell 112(2): 271–282.

Wyart, C., F. Del Bene, E. Warp, E. K. Scott, D. Trauner, H. Baier and E. Y. Isacoff (2009). “Optogenetic dissection of a behavioural module in the vertebrate spinal cord.” Nature 461(7262): 407–410.

Zimmerman, C. A., E. L. Huey, J. S. Ahn, L. R. Beutler, C. L. Tan, S. Kosar, L. Bai, Y. Chen, T. V. Corpuz, L. Madisen, H. Zeng and Z. A. Knight (2019). “A gut-to-brain signal of fluid osmolarity controls thirst satiation.” Nature 568(7750): 98–102.

